# Temperature causes species-specific responses to UV-induced DNA damage in amphibian larvae

**DOI:** 10.1101/2022.09.01.506274

**Authors:** Coen Hird, Craig E. Franklin, Rebecca L. Cramp

## Abstract

Anthropogenic ozone depletion has led to a 2-5% increase in ultraviolet B radiation (UVBR) levels reaching the earth’s surface. Exposure to UVBR causes harmful DNA damage in amphibians, but this is minimized by DNA repair enzymes such as thermally sensitive CPD-photolyase, with cool temperatures slowing repair rates. It is unknown whether amphibian species differ in the repair response to a given dose of UVBR across temperatures. We reared larvae of three species (*Limnodynastes peronii*, *Limnodynastes tasmaniensis*, and *Platyplectrum ornatum)* at 25°C and acutely exposed them to 80 μW cm^−2^ UVBR for 2 h at either 20°C or 30°C. UVBR-mediated DNA damage was measured as larvae repaired damage in photoreactive light at their exposure temperatures. Cool temperatures increased DNA damage in all two species and slowed DNA repair rate in *P. ornatum*. The magnitude of DNA damage incurred from UVBR was species-specific. *P. ornatum* had the lowest CPDs and DNA repair rates, and the depressive effects of low temperature on photorepair were greater in *L. tasmaniensis*. Considering the susceptibility of most aquatic organisms to UVBR, this research highlighted a need to consider the complexity of species-specific physiology when forecasting the influence of changing UVBR and temperature in aquatic ecosystems.

## INTRODUCTION

Solar ultraviolet (UV) radiation has wide-ranging effects on organisms and biological processes. Ultraviolet B radiation (UVBR) is largely absorbed by the stratospheric ozone column, however significant amounts of UVBR reach the Earth’s surface and penetrate aquatic ecosystems [1]. Anthropogenic O_3_ depletion led to a 2-5% increase in UVBR levels in some areas [2,3] and will likely remain high [4] or even possibly increase as a result of climate change [5–7]. UVBR can directly damage nucleic acids by forming dimers between nucleotide bases that disrupt gene expression patterns, cause mutations and trigger apoptosis [8,9]. To avoid the toxic effects of UV-induced DNA damage, organisms employ DNA repair mechanisms including nucleotide excision repair (NER) [10] and photoenzymatic repair (PER) [11]. During PER, the enzyme photolyase uses energy from photoreactive light to break UVBR-induced nucleotide bonds [12]. However, if DNA damage occurs at rates exceeding DNA repair, deleterious photoproducts may accumulate, resulting in whole-organism fitness consequences [13,14]. Aquatic organisms are susceptible to UVBR damage at the cellular level, leading to population impacts [15].

Globally, amphibians face a high extinction risk [16,17], [18]. Although a significant number of recent population declines have been linked to the emergence and spread of novel pathogenic amphibian chytrid fungi [19–22], global decline patterns point to co-occurring environmental changes, such as increasing UVBR, as proximate and interactive stressors with the pathogen [23–25]. Increased UVBR is hypothesised to influence amphibian populations through direct impacts on eggs and larvae as these life stages are often diurnal and typically laid during spring and summer when UVBR levels are highest [26]. UVBR exposure causes a range of sublethal and lethal effects in amphibian embryos and larvae [27] which occur primarily through the formation of cyclobutane pyrimidine photoproducts (CPDs) in DNA, which are repaired primarily through PER pathways in amphibians [28].

The negative effects of UVBR on amphibians are significantly compounded when UVBR exposure occurs at low temperatures [29–33], reflecting the thermal sensitivity of PER [34–39]. Morison et al. (2019) proposed that the thermal sensitivity of UVBR-associated DNA repair may partially explain why a disproportionately high number of amphibian declines have occurred at higher altitudes [25,40–46]. However, species from high UV or cool environments may differ in their tolerance to UVBR, by compensating for the depressive effect of temperature on DNA repair or by employing more efficient or effective DNA repair. In this study, we investigate the thermal sensitivity of UVBR effects on DNA in the larvae of three closely related amphibian species: *Limnodynastes peronii*, *Limnodynastes tasmaniensis*, and *Platyplectrum ornatum*. *L. peronii* larvae are known to be UVBR-sensitive [47–49] while *P. ornatum* larvae are considerably more UVBR tolerant and likely experience higher UVBR doses in nature [50]; the UVBR sensitivity of *L. tasmaniensis* larvae is unknown, however *L. tasmaniensis* are found in cooler habitats compared with *L. peronii*. It was hypothesised that there would be species-specific differences in the amount of incurred DNA damage, in DNA repair rates, and in the thermal dependence of DNA repair rates which reflect the differing thermal and UV environments inhabited by the three species.

## METHODS

### Animal collection and maintenance

Freshly laid *Limnodynastes peronii*, *Limnodynastes tasmaniensis*, and *Platyplectrum ornatum* spawn were collected near Meeanjin (greater Brisbane, QLD, Australia). Spawn was immediately transported to The University of Queensland, separated into small pieces and left to hatch in two litre ice-cream containers half-filled with carbon-filtered Brisbane tap water at 25°C. Partial water changes were conducted every second day and larvae were fed thawed spinach *ad libitum* daily. Larvae were reared for 2-3 weeks to Gosner stage 25 [51] under a 12L:12D photoperiod generated by standard room fluorescent lights.

### UVR lighting and heating

UVR was generated using 40 W, full spectrum fluorescent light sources which emit visible light, UVAR, and UVBR (Repti-Glo 10.0, 1200 mm, Exo Terra, Montreal, Canada). Light heights were adjusted to achieve an absolute UVBR irradiance of ~80 μW cm^−2^ (mean ± SD = 79.8 ± 4.9; UVAR irradiance: mean ± SD = 81.3 ± 54.5) at the water surface. Light intensities for UVAR and UVBR were measured using a calibrated radiometer (IL1400BL, International Light Inc., Newburyport, USA).

### Experimental design

Upon reaching Gosner stage 25, larvae were placed into individual wells of a six-well plate (34 plates per species) containing 10 ml of filtered water. The plates were evenly allocated across four water baths at 20°C or 30°C test temperatures (two per temperature). The temperature of experimental water baths was controlled using 300 W heaters (AquaOne, Kongs Pty Ltd, Ingleburn, NSW, Australia) and water was circulated by small pumps. Larvae were left for 1 h at the experimental temperature. Larvae (n = 72 in total, n=24 per species) were randomly removed from water baths (n=12 per temperature treatment) rapidly euthanised with buffered MS222 (0.25 mg L^−1^) and then snap frozen at −80°C. UVR lights were switched on and all remaining larvae were exposed to 80 μW cm^−2^ of UVBR for 2 h. Water baths were then covered with UVBR blocking film (Melinex 516, 100 μm, Archival Survival, Doncaster, Victoria, Australia) and larvae were allowed to photorepair for up to 24 h. Larvae (n = 72) were removed at the following time points post-UVBR exposure: 0, 0.25, 0.5, 0.75, 1, 1.5, 3, 6, 12, and 24 h), then euthanised and snap frozen. Larval wet mass was recorded (L. peronii mean ± SD = 5.7 ± 2; L. tasmaniensis mean ± SD = 9.5 ± 7.5; P. ornatum mean ± SD = 6.2 ± 3.1).

### DNA damage

Genomic DNA was extracted and purified from whole-animal homogenates using PureLink Genomic DNA Minikits (ThermoFisher Scientific Inc., Waltham, USA) and quantified using a Qubit dsDNA High-Range Assay Kit (ThermoFisher Scientific Inc., Waltham, MA, USA). Cyclobutane pyrimidine dimer (CPD) concentrations were determined using an anti-CPD ELISA assay following the primary antibody manufacturers protocol [52]. Briefly, DNA (0.4ng/uL) was loaded into triplicate wells of a protamine sulfate-coated 96-well plate and detected using an anti-CPD monoclonal primary antibody (NM-ND-D001, clone TDM-2, Cosmo Bio Co., Ltd.). The primary antibody was detected using a biotinylated goat anti-mouse IgG (QD209886, Life Technologies, USA), then an HRP-conjugated streptavidin (ab7403, Abcam, Cambridge, UK). Colour development was achieved with TMB substrate (416 mM; Sigma-Aldrich, Saint Louis, MO, USA) following the manufacturers guidelines. Colour development was stopped with H_2_SO_4_, and absorbance determined at 450 nm (Beckman Coulter DTX880 multimode detector, MN, USA) using the SoftMax® Pro program (Version 7.1.0, Molecular Devices LLC, CA, USA). CPD concentrations were calculated from a standard dose-response curve of UVC-irradiated calf thymus (NM-MA-R010, Cosmo Bio Co., Ltd., Tokyo, Japan) on each plate. CPD concentration is reported as units of UVCR-dose equivalent per 20 ng of DNA.

### Statistical analyses

All analyses were conducted in the R statistical environment [53]. CPD levels and wet body mass were log transformed to meet the assumptions of statistical tests. The decline in CPD abundance over time was interpreted as the repair rate of DNA damage. CPD abundance values >10 J m^2^ in a small number of larvae (n=11) were changed to 10 J m^2^ to ensure that these data fit within the dynamic range of the standard curve (0 – 10 J m^2^). ELISA Plate ID was recorded as a random effect. CPDs were fitted in a linear mixed effects model with the lme4 package [54] as:

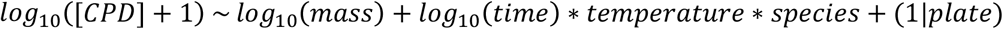

Where [CPD] was the UVCR-dose equivalent at three levels of species across 14 experimental ELISA 96-well plates, mass was the wet body mass of the tadpole in mg, and time was the hours post UVBR-exposure. The fitted model was analysed using omnibus Type II chi-square tests from the car package [55]. Specific contrasts between groups were generated from the lme4 package. Mean larval masses were compared between species using two-sample Wilcoxon rank sum tests with continuity correction. One-sample t-tests were used to confirm that pre-UVBR exposure (i.e., baseline) levels of CPDs were not significantly different from zero and whether CPD levels in 24 h post-exposure groups had returned to baseline levels for each species.

## RESULTS

Larvae that received no UVBR exposure had no CPDs (*L. peronii*: t_11_ = 1.48, *p* = 0.17; *L. tasmaniensis*: t_13_ = 1.89, *p* = 0.08; *P. ornatum*: t_14_ = 1.84, *p* = 0.09). For all species and temperatures, acute exposure to UVBR resulted in the formation of CPDs (Figure 1; Table S1; Table S2) which decreased over the 24 h recovery period but did not return to baseline (figure S1; *L. peronii*: t_13_ = 4.68, *p* <0.001; *L. tasmaniensis*: t_10_ = 3.51, *p* <0.01 ; *P. ornatum*: t_15_ = 4.35, *p* <0.001). Specific contrasts between regression coefficients were compared by cycling through three reference species in the full model (Table S2). There was no significant effect of temperature on DNA repair rates, but *L. tasmaniensis* at 20°C had higher rates of repair than *P. ornatum* at 20°C (Figure 1). There was a species-specific effect of temperature on CPD abundance (Figure 1). Larvae held at 20°C accumulated more CPDs than larvae held at 30°C in both *L. tasmaniensis* and *P. ornatum*, but not *L. peronii*. There were significant interspecific differences between CPD levels, dependent on the exposure temperature. At 20°C, *L. tasmaniensis* had higher CPDs than *P. ornatum*. At 30°C, *L. tasmaniensis* had lower CPDs than *L. peronii*, and *L. peronii* had higher CPDs than *P. ornatum*. Mean larval masses were significantly higher in L. tasmaniensis compared with L. peronii (W = 20537, *p* <0.001) and P. ornatum (W = 14012, *p* <0.001). P. ornatum larval mass was not significantly different to P. ornatum (W = 20263, *p* = 0.78).

**Figure 1.**
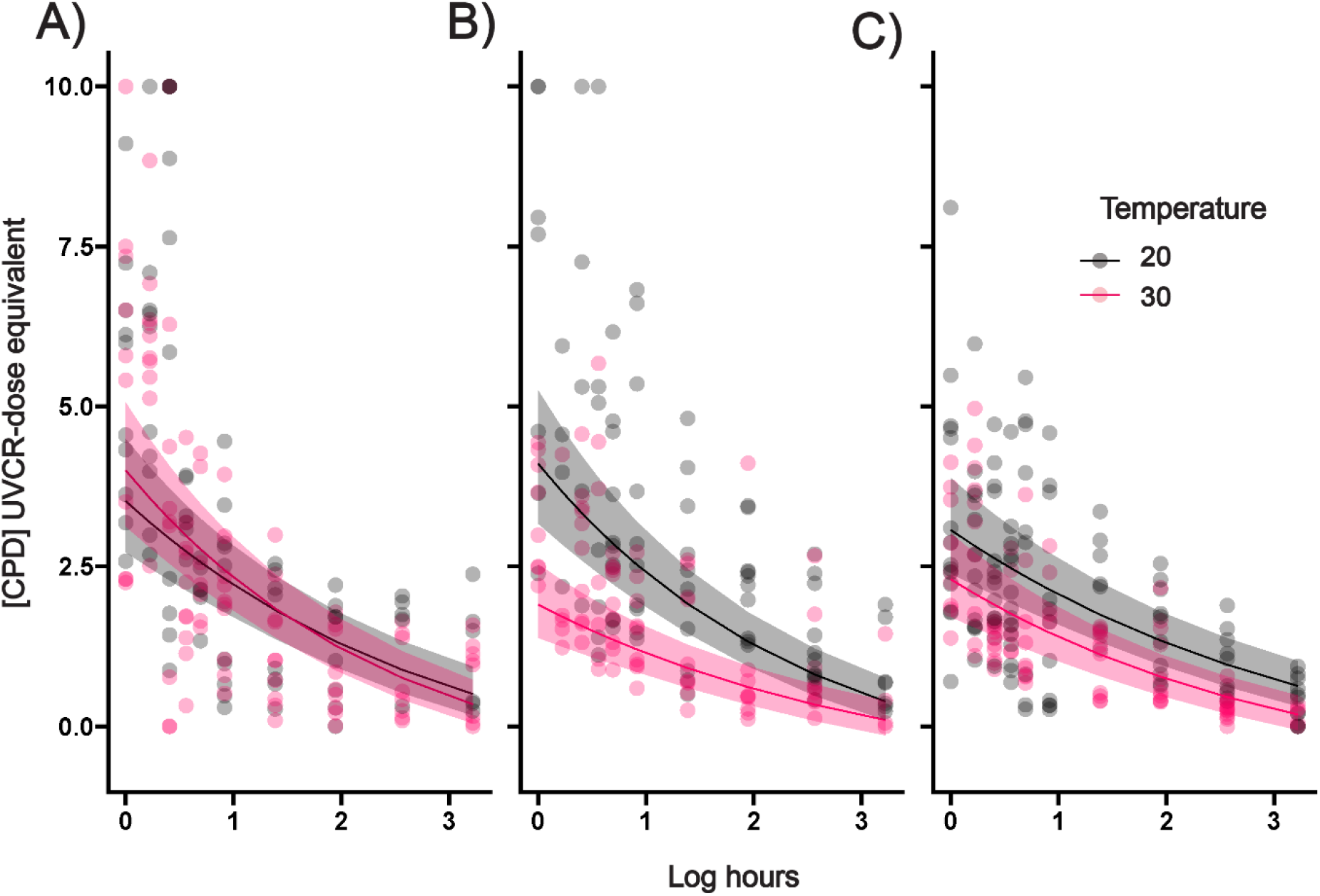
The effect of water temperature on CPD repair rates (measured as [CPD] UVCR-dose equivalent (J m^2^)) following 2 h of acute UVBR exposure (80 μw cm^−2^) and 24 h of blue light-assisted photorepair in A) *L. peronii*; B) *L. tasmaniensis*; and C) *P. ornatum* larvae. Points represent individual values. Curves represent the fitted parameters extracted from the linear mixed effects model, predicted for an average-sized tadpole of all species (6.08 mg). Ribbons show the upper and lower S.D. for the fitted line.

## DISCUSSION

This study demonstrated a complex interplay between temperature, and time on the levels of UV-induced DNA damage among three species of amphibian larvae. UVBR exposure caused significant DNA damage in all larvae, but there were considerable interspecific differences in the magnitude and thermal sensitivity of the damage incurred. At cool temperatures, *P. ornatum* accumulated less damage during UVBR exposure but had a lower rate of repair compared to *L. tasmaniensis*. Considering *P. ornatum* larvae likely experience greater natural UVBR levels than *L. peronii* and *L. tasmaniensis*, our data suggest that this species may use alternate strategies to prevent DNA damage from occurring, however, any that does occur is repaired more slowly than in the other two species. *P. ornatum* larvae can maintain performance over a relatively large range of environmental temperatures and in the presence of elevated UVBR [56]. Preventing DNA damage in *P. ornatum* may reduce metabolic costs associated with repair mechanisms. Metabolism underlies important fitness traits by providing energy for growth, development, and performance [57–59]. A lower DNA repair rate and lower amount of initial DNA damage following UVR exposure in *P. ornatum* may mean that more energy is available for other metabolic activities such as performance traits. However, the lower initial CPDs in *P. ornatum* does not necessarily mean the species is UV tolerant. Slower rates of DNA photorepair in *P. ornatum* may be problematic if larvae were to incur significant CPD concentrations in nature unless it also stimulated an increase in DNA repair rates.

Our study shows that cool temperatures did not lower UVBR-associated repair rates in any species, which contrasted with earlier work showing a thermal dependence of DNA repair rates in *L. peronii* [48]. This could indicate a concentration-dependent rate of photoenzymatic CPD repair, functioning antagonistically in cold environments with the depressive effects of temperature on enzyme activity. In the current experiment, larvae were subject to 80 μW cm^−2^ UVBR for 2 h which resulted in substantially greater DNA damage levels compared to the 100 μW cm^−2^ for 1 h exposure used by Morison et al. (2019). The additional hour of exposure time in the present study may be where differential rates of repair are occurring, rather than during the period of photorepair where larvae were sampled. This could also explain why CPD levels were higher in *L. peronii* and *L. tasmaniensis* larvae held at cool temperatures compared with warm temperatures, even though no difference in DNA repair rate were reported between temperatures. Comparing DNA abundance and repair across UV dose, duration, and intensity treatments may help to elucidate these different responses [60].

The finding that *L. peronii* accumulated a greater abundance of CPDs than *P. ornatum* at 20°C suggested that *L. tasmaniensis* larvae are more susceptible to UVR exposure at cool temperatures. This result was surprising given that *L. tasmaniensis* can occur in cooler habitats than *P. ornatum* and *L. peronii* and might be expected to be more resilient against the depressive effects of temperature on physiological rate processes. However, the *L. tasmaniensis* populations sampled in this study were not from particularly cool climates and may possess a thermal phenotype that more closely resembles that of *L. peronii* and *P. ornatum*. Population-level differences in thermal sensitivity can be underpinned by genotypic differences that may moderate the effects of temperature on the DNA photorepair response [61]. Alternatively, species-specific variations in cutaneous melanin concentration may explain differences in the degree of DNA damage experienced by larvae. Melanin pigments can migrate rapidly to the outer epithelium with UV insult to guard against DNA damage [62–64]. Differences in inherent melanin levels, or the capacity to rapidly mobilise melanin stores to minimise DNA damage could vary across species and with temperature. Similarly, sampling across a larger range of species may be useful to elucidate the contribution of phylogeny to the DNA damage response. Furthermore, there is evidence of thermal phenotypic plasticity in the DNA damage response in plants [65], but it is unknown whether amphibians possess the same. To determine whether larvae have the capacity for thermal phenotypic plasticity of the DNA photorepair response, larvae could be chronically reared at cooler and warmer temperatures prior to UV exposure.

Amphibians are among the world’s most threatened taxa, with global declines linked with co-occurring and interacting stressors such as disease and climate. Therefore, knowledge of how amphibians respond to environmental change is vital to their conservation. Our results show that UVR tolerance in amphibian larvae may depend upon the thermal context of their environment, but this influence depends on the species in question. Species and populations in which larvae can be more at risk of the interplay between harmful UVR exposure and cool temperatures may possess physiological adaptations enabling them to persist. However, these physiological differences could contribute to the differential susceptibility of species to decline, which may explain why a disproportionately high number of amphibian declines have occurred at high altitude. We argue that when attempting to predict how changing UVR and temperature levels in aquatic ecosystems has and will continue to influence amphibian larvae, it is critical to consider species-specific physiological responses.

## ACKNOWLEDGEMENTS

This research received support from an Australian Research Council Discovery grant awarded to CEF (DP190102152).

## ETHICS

Spawn was collected under the Queensland Department of Environment and Heritage Protection Scientific Purposes Permit (WA0017092) and procedures were approved by The University of Queensland’s Animal Ethics Unit (SBS/428/19).

## DATA AVAILABILITY STATEMENT

The complete datasets and R scripts used for analysing the data are publicly available at UQ eSpace (https://doi.org/10.48610/4fdfc26).

## AUTHOR CONTRIBUTIONS

Conceptualisation: RLC, CH, CEF; Methodology: CH, RLC, CEF; Validation: CH; Formal analysis: CH; Investigation: CH; Resources: CEF; Data curation: CH; Writing – original draft: CH; Writing – review & editing: CH, RLC, CEF; Visualisation: CH; Supervision: RLC, CEF; Project administration: RLC, CEF; Funding acquisition: RLC, CEF.

