## Supplemental file for "Temperature causes species-specific responses to UV-induced DNA damage in amphibian larvae"

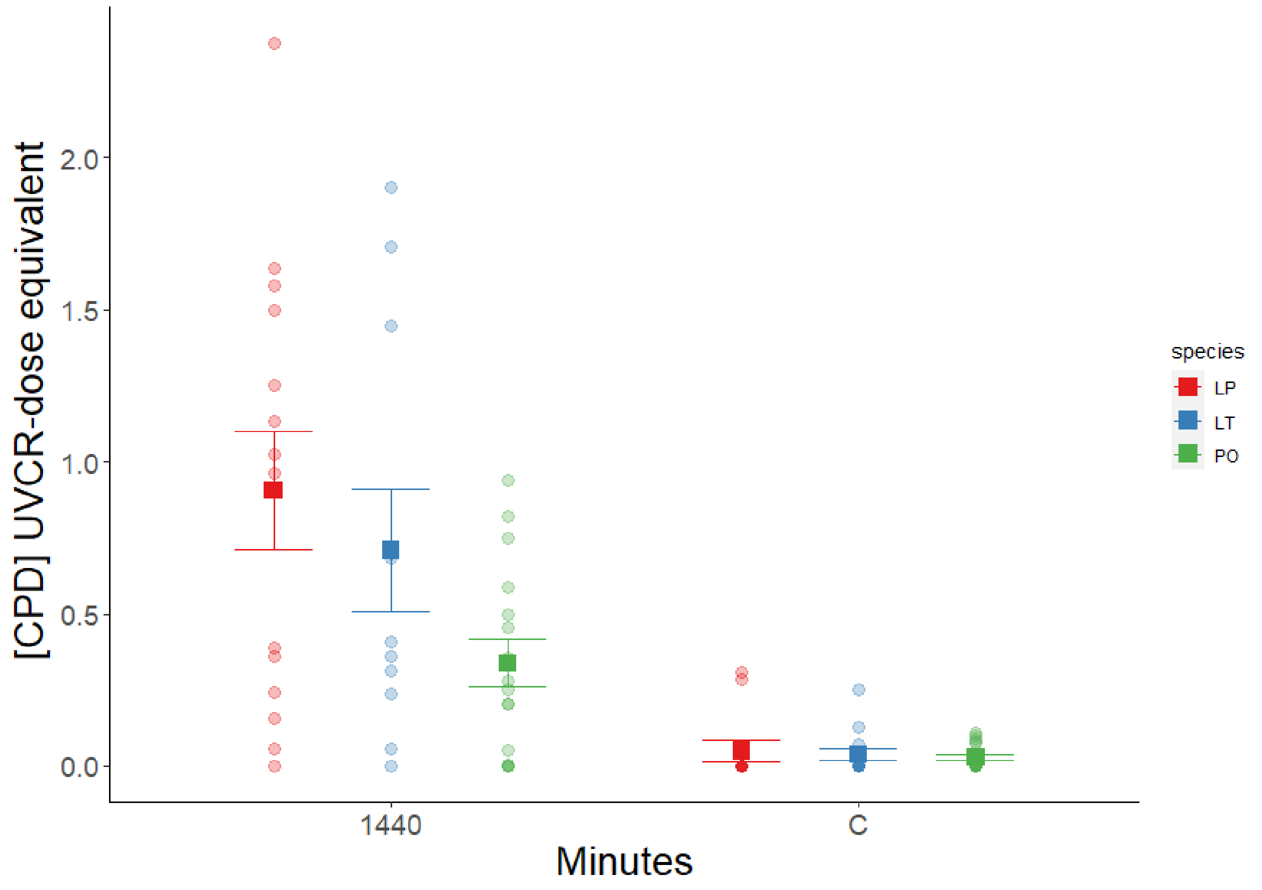


**Figure S1.** 24 hours of repair following 2 hours of acute UVBR exposure (80 μw $cm^{-2}$) does not result in the return of CPD levels to pre-exposure (control) concentrations in all species. Faint points represent raw data. Squares represent mean values and error bars represent standard error.

**Table S1.** Comparison of [CPD] following UVBR exposure in three species of amphibian larvae left to photorepair at two temperatures. Table generated using omnibus Type II chi-square tests from the car package.

| ***Predictors*** | ***X2*** | ***df*** | ***p*** |
|---|---|---|---|
| Mass | 15.3889 | 1 | **<0.001** |
| Time | 378.9278 | 1 | **<0.001** |
| Temperature | 43.2281 | 1 | **<0.001** |
| Species | 29.4556 | 2 | **<0.001** |
| Time:Temperature | 0.0318 | 1 | 0.85848 |
| Time:Species | 4.0044 | 2 | 0.13504 |
| Temperature:Species | 33.6339 | 2 | **<0.001** |
| Time:Temperature:Species | 5.3094 | 2 | 0.07032 |

**Table S2.** Contrasts of [CPD] between coefficients following UVBR exposure in three species of amphibian larvae left to photorepair at two temperatures. Table generated with the sjPlot package from the linear mixed effects model generated from the lme4 package. Separate contrasts are presented using all combinations of species and temperatures as the reference group for contrasts.

| Reference group | Predictors | Estimates | CI | p | df |
|---|---|---|---|---|---|
| ***L. peronii* (20°C)** | (Intercept) | 1.72 | 1.50 – 1.94 | **<0.001** | 46.62 |
| | logmass mg | -0.12 | -0.18 – -0.06 | **<0.001** | 510.61 |
| | logtime | -0.34 | -0.43 – -0.26 | **<0.001** | 516.46 |
| | temp [30] | 0.1 | -0.06 – 0.26 | 0.216 | 509.5 |
| | species [LT] | 0.12 | -0.06 – 0.31 | 0.194 | 516.96 |
| | species [PO] | -0.1 | -0.26 – 0.05 | 0.181 | 512.21 |
| | logtime * temp [30] | -0.07 | -0.17 – 0.04 | 0.226 | 509.9 |
| | logtime * species [LT] | -0.06 | -0.18 – 0.06 | 0.296 | 516.71 |
| | logtime * species [PO] | 0.06 | -0.05 – 0.16 | 0.297 | 514.11 |
| | temp [30] * species [LT] | -0.67 | -0.91 – -0.42 | **<0.001** | 514.13 |
| | temp [30] * species [PO] | -0.31 | -0.54 – -0.08 | **0.009** | 513.64 |
| | (logtime * temp [30]) * species [LT] | 0.17 | 0.01 – 0.32 | **0.032** | 510.26 |
| | (logtime * temp [30]) * species [PO] | 0.03 | -0.11 – 0.18 | 0.665 | 510.65 |
| ***L. peronii* (30°C)** | (Intercept) | 1.82 | 1.60 – 2.04 | **<0.001** | 46.11 |
| | logmass mg | -0.12 | -0.18 – -0.06 | **<0.001** | 510.61 |
| | logtime | -0.41 | -0.49 – -0.32 | **<0.001** | 506.88 |
| | temp [20] | -0.1 | -0.26 – 0.06 | 0.216 | 509.5 |
| | species [LT] | -0.54 | -0.72 – -0.37 | **<0.001** | 515.84 |
| | species [PO] | -0.42 | -0.58 – -0.25 | **<0.001** | 514.25 |
| | logtime * temp [20] | 0.07 | -0.04 – 0.17 | 0.226 | 509.9 |
| | logtime * species [LT] | 0.11 | -0.01 – 0.23 | 0.08 | 516.59 |
| | logtime * species [PO] | 0.09 | -0.02 – 0.20 | 0.121 | 516.48 |
| | temp [20] * species [LT] | 0.67 | 0.42 – 0.91 | **<0.001** | 514.13 |

| | | | | | |
|---|---|---|---|---|---|
| | temp [20] * species [PO] | 0.31 | 0.08 – 0.54 | **0.009** | 513.64 |
| | (logtime * temp [20]) * species [LT] | -0.17 | -0.32 – -0.01 | **0.032** | 510.26 |
| | (logtime * temp [20]) * species [PO] | -0.03 | -0.18 – 0.11 | 0.665 | 510.65 |
| ***L. tasmaniensis* (20°C)** | (Intercept) | 1.84 | 1.60 – 2.09 | **<0.001** | 66.67 |
| | logmass mg | -0.12 | -0.18 – -0.06 | **<0.001** | 510.61 |
| | logtime | -0.4 | -0.49 – -0.32 | **<0.001** | 516.99 |
| | temp [30] | -0.56 | -0.75 – -0.38 | **<0.001** | 516.2 |
| | species [LP] | -0.12 | -0.31 – 0.06 | 0.194 | 516.96 |
| | species [PO] | -0.23 | -0.40 – -0.05 | **0.011** | 516.56 |
| | logtime * temp [30] | 0.1 | -0.01 – 0.21 | 0.07 | 510.36 |
| | logtime * species [LP] | 0.06 | -0.06 – 0.18 | 0.296 | 516.71 |
| | logtime * species [PO] | 0.12 | 0.02 – 0.23 | **0.025** | 515.57 |
| | temp [30] * species [LP] | 0.67 | 0.42 – 0.91 | **<0.001** | 514.13 |
| | temp [30] * species [PO] | 0.35 | 0.11 – 0.60 | **0.004** | 515.44 |
| | (logtime * temp [30]) * species [LP] | -0.17 | -0.32 – -0.01 | **0.032** | 510.26 |
| | (logtime * temp [30]) * species [PO] | -0.14 | -0.28 – 0.01 | 0.068 | 509.02 |
| ***L. tasmaniensis* (30°C)** | (Intercept) | 1.28 | 1.05 – 1.51 | **<0.001** | 57.8 |
| | logmass mg | -0.12 | -0.18 – -0.06 | **<0.001** | 510.61 |
| | logtime | -0.3 | -0.38 – -0.22 | **<0.001** | 512.45 |
| | temp [20] | 0.56 | 0.38 – 0.75 | **<0.001** | 516.2 |
| | species [LP] | 0.54 | 0.37 – 0.72 | **<0.001** | 515.84 |
| | species [PO] | 0.13 | -0.05 – 0.31 | 0.169 | 516.25 |
| | logtime * temp [20] | -0.1 | -0.21 – 0.01 | 0.07 | 510.36 |
| | logtime * species [LP] | -0.11 | -0.23 – 0.01 | 0.08 | 516.59 |
| | logtime * species [PO] | -0.02 | -0.13 – 0.09 | 0.768 | 510.38 |
| | temp [20] * species [LP] | -0.67 | -0.91 – -0.42 | **<0.001** | 514.13 |
| | temp [20] * species [PO] | -0.35 | -0.60 – -0.11 | **0.004** | 515.44 |
| | (logtime * temp [20]) * species [LP] | 0.17 | 0.01 – 0.32 | **0.032** | 510.26 |
| | (logtime * temp [20]) * species [PO] | 0.14 | -0.01 – 0.28 | 0.068 | 509.02 |
| ***P. ornatum* (20°C)** | (Intercept) | 1.62 | 1.41 – 1.83 | **<0.001** | 37.55 |

| | | | | | |
|---|---|---|---|---|---|
| | logmass mg | -0.12 | -0.18 – -0.06 | **<0.001** | 510.61 |
| | logtime | -0.28 | -0.35 – -0.22 | **<0.001** | 508.17 |
| | temp [30] | -0.21 | -0.37 – -0.05 | **0.01** | 514.48 |
| | species [LP] | 0.1 | -0.05 – 0.26 | 0.181 | 512.21 |
| | species [LT] | 0.23 | 0.05 – 0.40 | **0.011** | 516.56 |
| | logtime * temp [30] | -0.03 | -0.13 – 0.06 | 0.488 | 507.62 |
| | logtime * species [LP] | -0.06 | -0.16 – 0.05 | 0.297 | 514.11 |
| | logtime * species [LT] | -0.12 | -0.23 – -0.02 | **0.025** | 515.57 |
| | temp [30] * species [LP] | 0.31 | 0.08 – 0.54 | **0.009** | 513.64 |
| | temp [30] * species [LT] | -0.35 | -0.60 – -0.11 | **0.004** | 515.44 |
| | (logtime * temp [30]) * species [LP] | -0.03 | -0.18 – 0.11 | 0.665 | 510.65 |
| | (logtime * temp [30]) * species [LT] | 0.14 | -0.01 – 0.28 | 0.068 | 509.02 |
| **P. ornatum (30°C)** | (Intercept) | 1.41 | 1.19 – 1.62 | **<0.001** | 43.22 |
| | logmass mg | -0.12 | -0.18 – -0.06 | **<0.001** | 510.61 |
| | logtime | -0.32 | -0.39 – -0.24 | **<0.001** | 507.34 |
| | temp [20] | 0.21 | 0.05 – 0.37 | **0.01** | 514.48 |
| | species [LP] | 0.42 | 0.25 – 0.58 | **<0.001** | 514.25 |
| | species [LT] | -0.13 | -0.31 – 0.05 | 0.169 | 516.25 |
| | logtime * temp [20] | 0.03 | -0.06 – 0.13 | 0.488 | 507.62 |
| | logtime * species [LP] | -0.09 | -0.20 – 0.02 | 0.121 | 516.48 |
| | logtime * species [LT] | 0.02 | -0.09 – 0.13 | 0.768 | 510.38 |
| | temp [20] * species [LP] | -0.31 | -0.54 – -0.08 | **0.009** | 513.64 |
| | temp [20] * species [LT] | 0.35 | 0.11 – 0.60 | **0.004** | 515.44 |
| | (logtime * temp [20]) * species [LP] | 0.03 | -0.11 – 0.18 | 0.665 | 510.65 |
| | (logtime * temp [20]) * species [LT] | -0.14 | -0.28 – 0.01 | 0.068 | 509.02 |
